# Identification, characterization and transcriptional analysis of the long non-coding RNA landscape in the family *Cucurbitaceae*

**DOI:** 10.1101/2024.01.12.575433

**Authors:** Pascual Villalba-Bermell, Joan Marquez-Molins, Gustavo Gomez

## Abstract

Long non-coding RNAs (lncRNAs) constitute a fascinating class of regulatory RNAs, widely distributed in eukaryotes. In plants, they exhibit features such as tissue-specific expression, spatiotemporal regulation, and responsiveness to stress, suggesting their involvement in specific biological processes. Although an increasing number of studies support the regulatory role of lncRNAs in model plants, our knowledge about these transcripts in relevant crops is limited. In this study we employ a custom pipeline on a dataset of over 1,000 RNA-seq studies across nine representative species of the family *Cucurbitaceae* to predict 91,209 non-redundant lncRNAs. LncRNAs were predicted according to three confidence levels and classified into intergenic, natural antisense, intronic, and sense overlapping. Predicted lncRNAs have lower expression levels compared to protein-coding genes but a more specific behavior when considering plant tissues, developmental stages, and response to stress, emphasizing their potential roles in regulating various aspects of plant-biology. The evolutionary analysis indicates higher positional conservation than sequence conservation, which may be linked to the presence of conserved modular motifs within syntenic lncRNAs. In short, this research provides a comprehensive map of lncRNAs in the agriculturally relevant *Cucurbitaceae* family, offering a valuable resource for future investigations in crop improvement.

## INTRODUCTION

Plants, as sessile organisms, have evolved intricate mechanisms to adapt to changing environmental conditions and regulate their development (1). Deciphering the complex regulatory networks that underlie these processes has been a long-standing challenge in plant biology. Traditionally, protein-coding (PC) genes have been the primary focus of research, but recent advancements in high-throughput sequencing and functional genomics have unveiled the vast landscape of non-coding RNAs (ncRNAs), including a group identified as long non-coding RNAs (lncRNAs), which were initially considered as transcriptional “noise” (2).

Long non-coding RNAs represent a diverse class of transcripts characterized by their length (>200 nucleotides) and lack of protein-coding potential. These molecules, that compared to PC genes are generally expressed at lower levels, exhibit distinct features, such as tissue-specific expression patterns, spatiotemporal regulation, expression related to environmental changes and subcellular localization, suggesting their involvement in specific biological processes (3).

Regarding their biogenesis, lncRNAs are usually transcribed by RNA polymerase (Pol) II and subjected to 5’ capping and 3’ polyadenilation (3, 4). Occasionally, plant lncRNAs can be also dependent of RNA Pol III, Pol IV and/or Pol V (5, 6). However, this non-canonical lncRNAs are poorly characterized due mainly to their low expression and high instability (7).

Based on their genomic origin, orientation, and proximity to protein-coding genes plant lncRNAs can be commonly classified in four main classes: i) intergenic lncRNAs (lincRNAs) that do not overlap with other genes, ii) sense lncRNA (SOT-lncRNAs) that overlap (total or partially) with the same strand of its associated PC gene, iii) antisense lncRNA (NAT-lncRNAs) that overlap (total or partially) with the opposite strand of its associated gene, and iv) intronic lncRNAs (int-lncRNAs), located within an intron of the associated PC gene. Both, NAT-lncRNAs and lincRNAs constitutes the predominant classes of lncRNAs described and characterized in plants (8).

Since the discovery in 1994 of ENOD40 (EARLY NODULIN 40) the first lncRNA described in plants (9) and with the advancement of sequencing technologies, an increasing number of lncRNAs have been identified and functionally validated in diverse crops and model plants (8). These studies have demonstrated that plant lncRNAs contribute to various aspects of plant biology, including development, stress responses, genome stability, photo-morphogenesis, reproduction, flowering, and stress responses (10). Under a functional viewpoint, lncRNAs exert their regulatory roles through diverse mechanisms, such as transcriptional regulation, chromatin remodeling, epigenetic modifications, RNA splicing, and post-transcriptional regulation (11). Although, functional studies of plant lncRNAs are at an early stage, it is currently accepted that the availability of a comprehensive atlas of lncRNAs in crop plants, will enable us to use lncRNAs as potential, biomarkers and/or traits, in breading for tailoring stress-tolerant plants (12, 13).

The *Cucurbitaceae* (cucurbits) constitutes an important family of plants including most of 900 species in over 90 genera that are mainly distributed in tropical and subtropical areas (14). Some cucurbits members, that were domesticated and subsequently cultivated for thousands of years, represent agronomical important crops, including cucumber (*Cucumis sativus*), melon (*Cucumis melo*), watermelon (*Citrullus lanatus*), pumpkin and squash (*Cucurbita pepo*, *C. moschata*, *C. maxima* and *C. agyrosperma*) and bitter gourd (*Momordica charantia*). Others are used as medicinal plants (*Citrullus colocynthis*) or have practical uses for example as a bottle (*Lagenaria siceraria*) (15). In 2020, the area cultivated with cucurbits worldwide was 10.42 million hectares with a yielded near to 360 million tons (http://faostat.fao.org). Furthermore, members of the *Cucurbitaceae* family have been extensively used as model organisms to study fundamental biological processes such as vascular development (16), RNA trafficking (17), sex determination (18) fruit ripening (19) and host epigenetic alterations associated to infection (20). Consequently, deciphering the molecular mechanisms underlying cucurbit biology is of significant importance for improving crop yield, quality, and resilience in response to changing environments (21).

Recent studies have revealed the potential importance of lncRNAs as modulators of the development (22, 23) or the response to biotic (24–26) or abiotic (27, 28) stress conditions in diverse *Cucurbitaceae* family members. However, detailed information about lncRNAs in cucurbits is limited, and only a list of unclassified and barely characterized lncRNAs in cucumber, melon and watermelon are currently available in GreeNC (29), PLncDB V2.0 (30) or CANTATAdb 2.0 (31), the most relevant databases containing plant lncRNAs. Notably, no standardized information about lncRNAs can be found in the Cucurbit Genomics Database (32), that is the reference portal for the genomic annotation of members of this family. Thus revealing the lack of studies able to offer detailed information about the global landscape of these regulatory RNAs in cucurbits.

Here, we use a vast dataset of more than 1000 RNA-seq studies to gain insights into the identity, characteristics and expression of lncRNAs in nine representative species of the family *Cucurbitaceae*. Using a custom pipeline we have identified intergenic, natural antisense, intronic and sense overlapping lncRNAs, finding that distinct molecular features are associated to each type. Finally we have analyzed the evolutionary relationships and the expression in different tissues, developmental stages and stress conditions of the predicted lncRNAs. Overall, this exhaustive study reveals the diversity and molecular features of these emerging regulatory players in cucurbits, providing the first comprehensive map of lncRNAs in this agronomical relevant family.

## RESULTS

### Strategy for lncRNA prediction

A graphical resume of the pipeline developed in this work for identification, classification and characterization of lncRNAs in cucurbits is depicted in Figure 1. All the available RNA-seq data corresponding to 12 cucurbit species comprising 3494 RNA-seq libraries and 271 RNA-seq projects were downloaded from SRA database of the NCBI (https://www.ncbi.nlm.nih.gov/sra), the corresponding SRA accession codes, quality control and strandedness information are detailed in the Table S1. Three species having only one strand-specific RNA-seq sample (*B. hispida, L. aegyptiaca and S. grosvenorii*) were removed from our dataset. Consequently, after quality control and strandedness inference, 9 species (*C. sativus, C. melo, C. lanatus, L. siceraria, C. moschata, C. argyrosperma, C. pepo, C. maxima and M. charantia*) comprising 1116 cleaned and strand-specific RNA-seq libraries and 78 RNA-seq projects were included in the subsequent analysis (Table 1). The nine analized species constitute representative members of three different tribes and five genus in the *Cucurbitaceae* family (14).

**Figure 1.**
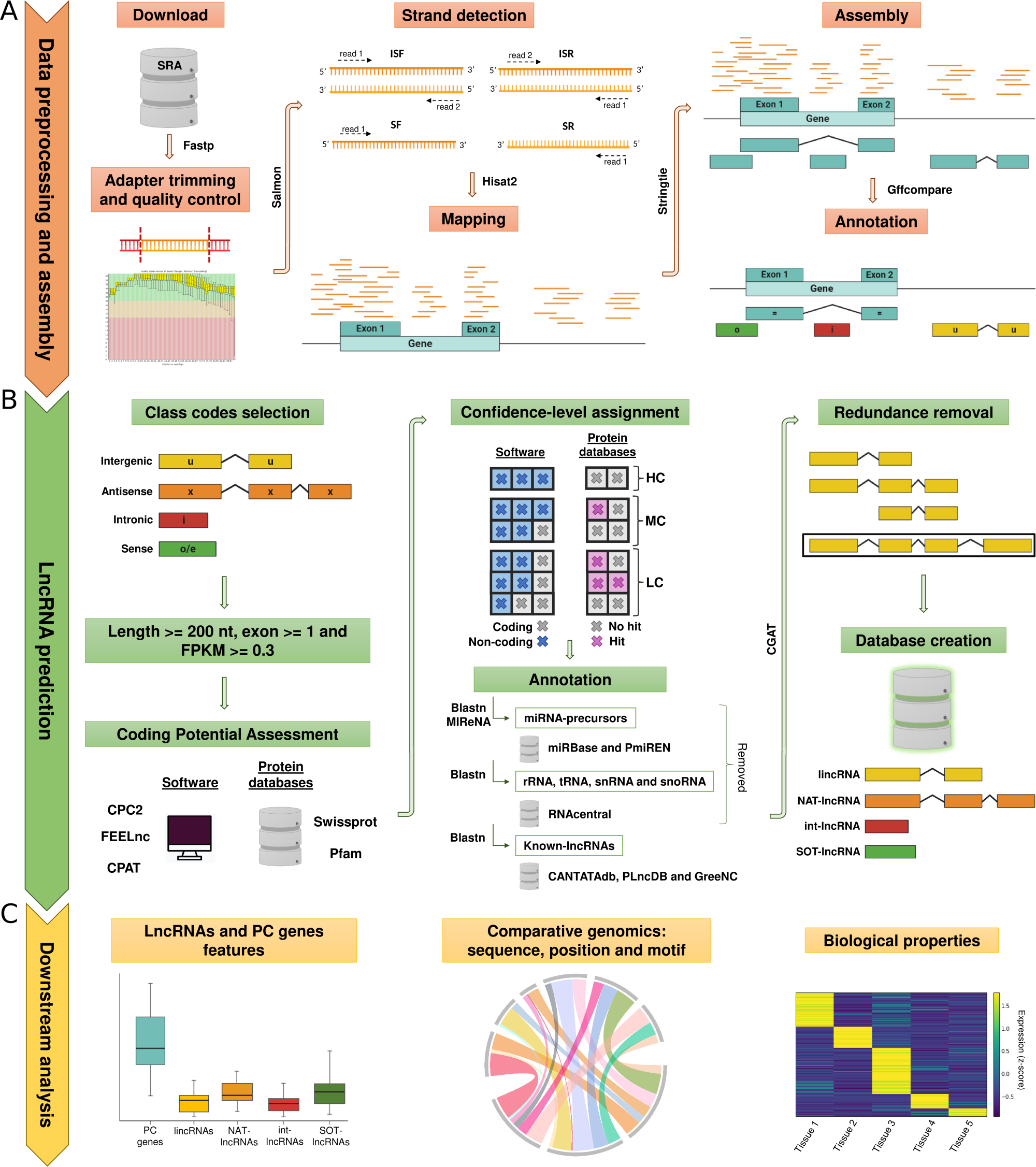
Graphic representation of the bioinformatic workflow used for the prediction, classification and analysis of lncRNAs. (A) Data recovering, preprocessing and transcriptome assembly. (B) Prediction and categorization of lncRNAs: intergenic (lincRNAs), natural antisense (NAT-lncRNAs), intronic (int-lncRNAs) and sense overlapping (SOT-lncRNAs) lncRNAs. (C) Downstream analysis to compare lncRNA features, conservation at three levels (sequence, genomic position and motifs) and differential expression (related to tissue, development and environment).

**Table 1:** Information of the analyzed datasets.

| Species | ID | Genome Size<br>(bases; Mb) | Samples (number) |  |  | Projects (number) |  |  | Final Data Size<br>(bytes; Gb) |
| --- | --- | --- | --- | --- | --- | --- | --- | --- | --- |
|  |  |  | Download | Trimming<br>And QC | Strand<br>Specific | Download | Trimming<br>And QC | Strand<br>Specific |  |
| *Cucumis sativus* | csa | 226.21 | 1388 | 1334 | 360 | 129 | 127 | 35 | 1167.43 |
| *Cucumis melo* | cme | 357.86 | 802 | 776 | 383 | 45 | 44 | 16 | 820.87 |
| *Citrullus lanatus* | cla | 365.45 | 717 | 711 | 231 | 55 | 54 | 17 | 663.61 |
| *Lagenaria siceraria* | lsi | 313.81 | 92 | 92 | 9 | 7 | 7 | 3 | 27.07 |
| *Cucurbita moschata* | cmo | 273.42 | 127 | 126 | 39 | 17 | 16 | 6 | 102.73 |
| *Cucurbita argyrosperma* | car | 231.58 | 10 | 10 | 9 | 2 | 2 | 2 | 30.36 |
| *Cucurbita pepo* | cpe | 263.38 | 143 | 142 | 50 | 13 | 13 | 7 | 112.61 |
| *Cucurbita maxima* | cma | 279.69 | 50 | 50 | 27 | 10 | 10 | 4 | 43.36 |
| *Momordica charantia* | mch | 294.01 | 74 | 74 | 8 | 5 | 5 | 2 | 27.73 |

Cleaned reads were mapped to the corresponding reference genome and assembled to create a merged transcriptome for each analyzed species (Figure 1A and “Material and Methods”). Comparable levels of genome coverage were observed across the selected species (Figure S1A). Assembled transcripts were categorized according to their position in the genome relative to protein-coding (PC) genes and those classified as intergenic (“u”), antisense (“x”), intronic (“i”) and sense (“o” or “e”) were selected. Only non-redundant transcripts, longer than 200 nucleotides and expression level above 0.3 Fragments Per Kilobase Million (FPKM) in at least one of the experiments, were considered. As detailed in Material and methods, we used three tools: CPC2 (33), FEELnc (34) and CPAT (35) to predict lncRNAs. Moreover, we looked for homology with known proteins and domains in SwissProt and Pfam databases, respectively. It is important to remark that our strategy for lncRNA prediction was designed to keep a balance between sensitivity and robustness. Consequently, inferred lncRNAs were classified into three confidence levels, according to the fulfillment of the following criteria: i) *High* (HC): predicted as lncRNA by the three software (CPC2, CPAT and FEELnc) and lacking homology with ORFs in the protein databases, ii) *Medium* (MC): predicted as lncRNA by the three software but have similarity with one of the protein databases or lack similarities with the two protein databases but are only predicted by two of the three software and iii) *Low* (LC): predicted as lncRNA by two of the software or only one software and lack similarity with one of the protein databases.

Once excluded potential lncRNAs homologous to miRNAs precursors, ribosomal, transfer, small nuclear and small nucleolar RNAs (see “Material and Methods” for details), transcripts were aligned and compared against those deposited in three plant lncRNAs databases (PLncDB V2.0, CANTATAdb 2.0 and GreeNC). The predicted lncRNAs were classified according to their genomic location into intergenic lncRNAs (lincRNAs), natural antisense lncRNAs (NAT-lncRNAs), intronic lncRNAs (int-lncRNAs) and sense overlapping lncRNAs (SOT-lncRNAs). Finally, further analyses were performed to compare the characteristics, conservation and expression profiles of the identified lncRNAs (Figure 1C).

### A wide range of lncRNA are distributed through cucurbit genomes

Following the strategy previously described, we were able to predict 91209 non-redundant lncRNAs in the analyzed cucurbits (Figure 2A and Table S2). The highest number of potential lncRNAs (29140) was identified in *C. melo*. Comparable amounts of lncRNAs were described in *C. sativus* (13563), *C. lanatus* (11189) and *C. argyrosperma* (10077). While *C. pepo* (8172), *C. moschata* (5472), *C. maxima* (4672), *M. charantia* (4515) and *L. siceraria* (4409), were the species with less predicted lncRNAs (Table S2). The lncRNAs categorized as High Confidence (HC), were the predominant class identified in the nine analyzed cucurbits (with percentages ranging from 68% to 84%). In contrast, those classified as Low Confidence (LC) were the less abundant (4.5% to 12.4%). Intermediate values (9.3% to 19,3%) were obtained for lncRNAs categorized as Medium Confidence (MC) (Table S2).

**Figure 2.**
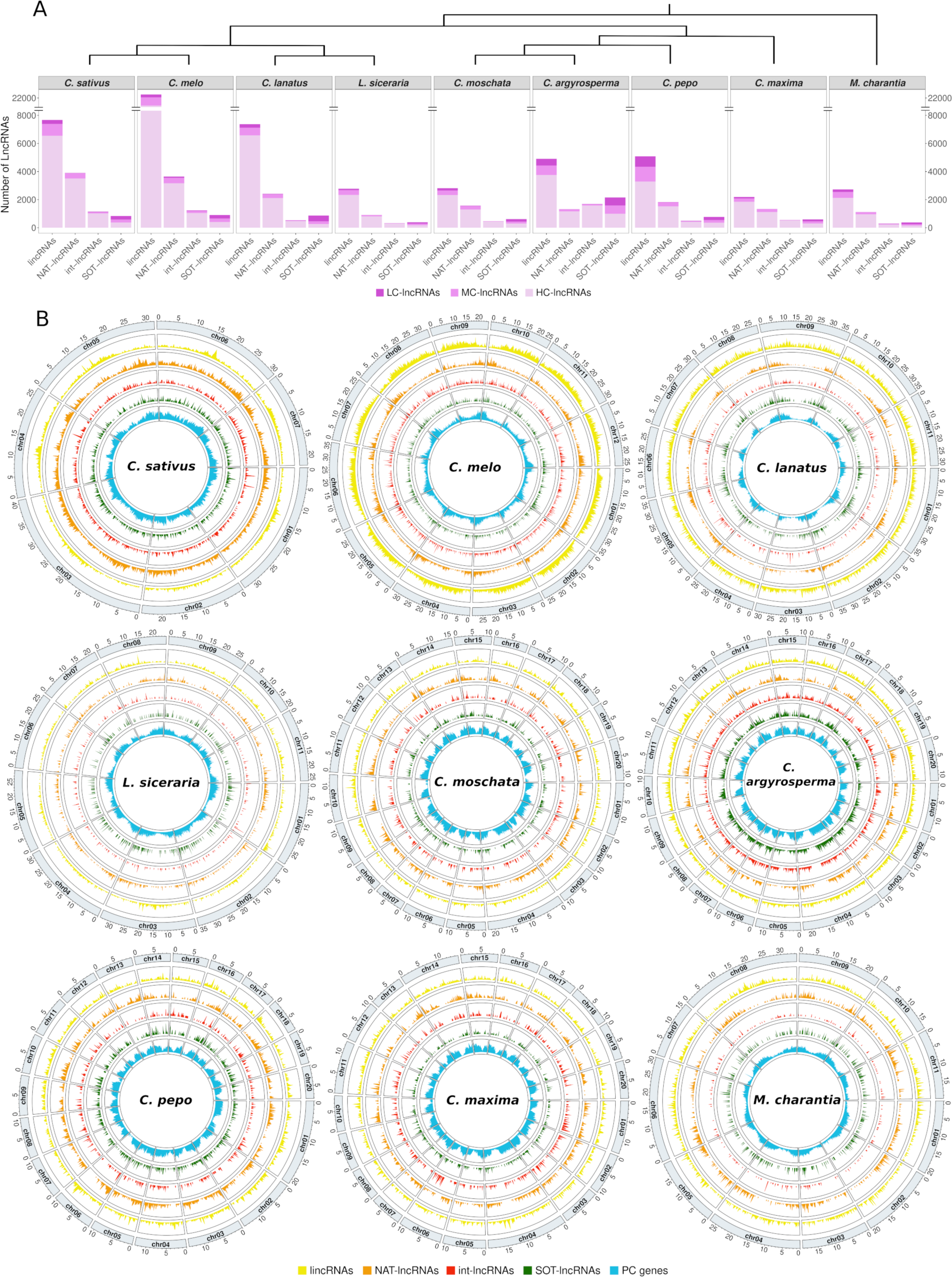
Detailed description and global landscape of the predicted lncRNAs in cucurbits. (A) Bar plot representing the number of lncRNAs identified in the nine species analyzed of the family *Cucurbitaceae*. These are classified into intergenic (lincRNAs), natural antisense (NAT-lncRNAs), intronic (int-lncRNAs) and sense overlapping (SOT-lncRNAs) lncRNAs, and categorized according to the confidence level of the prediction into: High-Confidence (HC-lncRNAs), Medium-Confidence (MC-lncRNAs) and Low-Confidence (LC-lncRNAs). The schematic tree on upper part of the plot indicates phylogenetic relationships among the species (1). (B) Circos plots representing the genomic distribution of the four types of the identified lncRNAs at the three confidence levels and annotated protein-coding genes (PC genes) in the nine species analyzed. Density is measured as the number of transcripts by window using a 250Kb window size.

According to their genomic context, intergenic lncRNAs (lincRNAs) were the most abundant type in the nine analyzed species, with 58907 potential lncRNAs representing the 64.5% of the total. They were followed by those classified as natural antisense lncRNAs (NAT-lncRNAs), which with a total number of 18050 lncRNAs were (except in *C. argyrosperma*) the second most abundant class (Table S2). The sense overlapping lncRNAs (SOT-lncRNAs) and intronic lncRNAs (int-lncRNAs) with 6742 and 7510 identified transcripts respectively, were the less represented and their prevalence varied depending on the species (Table S2).

In order to study the genomic distribution of these regulatory RNAs, we compared the localization pattern of the different types of lncRNAs with that of PC genes. As it is showed in the Figure 2B, the predicted lncRNAs were distributed throughout the genomes of all nine analyzed cucurbits and did not show such a clear pattern as the predominant localization of PC genes in non-centromeric regions. The mean genome covered by the predicted lncRNAs (5.13%) was considerably lower than the observed for PC genes (35.86%) (Table S3 and Figure S1B and C). Considering the categorization by the prediction confidence, we observed that the HC-lncRNAs cover a higher percentage of genome (3.73%) than MC-lncRNA (0.76%) and LC-lncRNAs (0.64%) (Table S3 and Figure S1B and C). The specie-specific analysis showed that the HC-lncRNAs predicted in *C. sativus* and *C. melo* exhibit the highest values of genome covered (7.60% and 7.84%, respectively) of the nine cucurbits.

To have a more quantitative perspective about the spatial distribution, we defined high-density regions (see “Material and Methods”) for each type of lncRNA and also for PC genes in the nine species and estimated their overlap (Figure S2). Our results show that the major overlap values of high-density regions was observed between NAT-lncRNAs and PC genes (median of 31.53%), while values below the 9% were observed for the remaining lncRNA types (Figure S2A). Comparable results were obtained when the numbers of overlapping high-density regions were individually determined in the nine cucurbits species (Figure S2B).

### Predicted lncRNAs show significant sequence differences with coding-RNAs

To provide a more detailed view about their molecular properties, we analyzed the GC content, exon number, length, expression and content of repetitive sequences of the predicted lncRNAs, using as references PC genes and a random intergenic region (when this comparison was possible, see “Material and Methods” for more details). In general, the values observed in lncRNAs were significantly lower than those in PC genes, except for the content of repetitive regions where the value obtained for lncRNAs was significantly higher (Figure 3A and Table S4). The most evident difference was found for the expression level estimated as Transcript Per Million (TPM), where the lncRNAs (median of 0.40 TPM) were expressed about 20-fold lower levels than PC genes (median of 8.21 TPM). Considering the differences with intergenic regions, while lncRNAs showed a significantly higher GC content (median of 39.02% and 34.80% for lncRNAs and intergenic regions, respectively), no significant difference was found in the content of repetitive sequences. It is worthy to note that similar differences were observed when this comparison was individually performed for each one of the nine cucurbits species (Figure S3 and Table S4).

**Figure 3.**
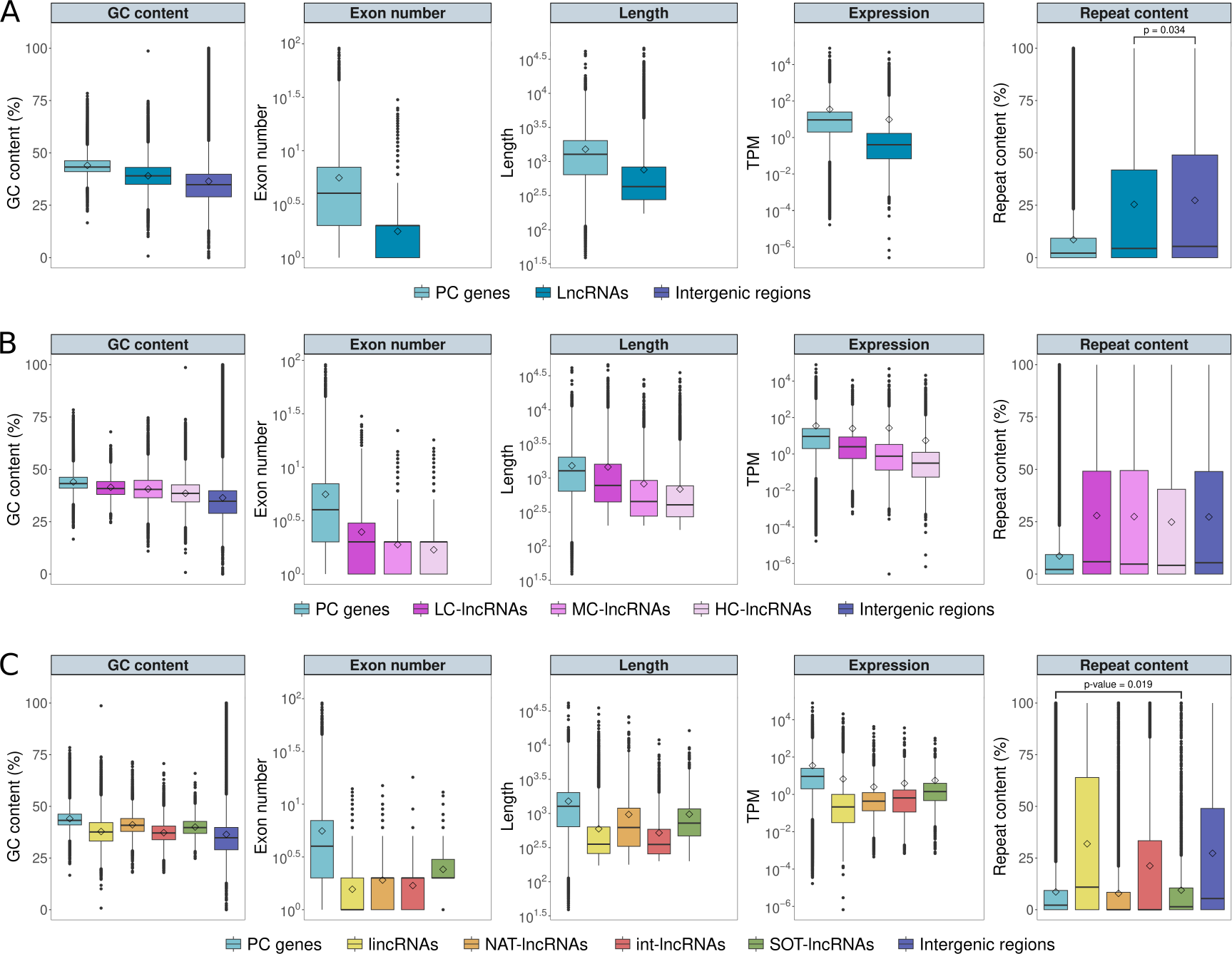
Molecular properties of the identified lncRNAs of cucurbits. Box plots showing the comparisons of GC content, number of exons, transcript length, expression levels and repetitive content features between lncRNAs, protein-coding genes (PC genes) and random intergenic regions (when corresponding): (A) Considering all the identified lncRNAs, (B) Detailed information of the lncRNAs separated according to the confidence level of the prediction: High-Confidence (HC-lncRNAs), Medium-Confidence (MC-lncRNAs) and Low-Confidence (LC-lncRNAs), (C) HC-lncRNAs separated in the four classes: intergenic (lincRNAs), natural antisense (NAT-lncRNAs), intronic (int-lncRNAs) and sense overlapping (SOT-lncRNAs) lncRNAs. Differences between pairs of box plots within each facet are statistically significant (Wilcoxon rank sum test p-value < 0.01), unless indicated otherwise. In each boxplot the mean value is represented by a square. Information on box colors is detailed below each panel.

Finally, considering the confidence of the lncRNA prediction, the observed differences with PC genes and intergenic regions were more evident in lncRNAs classified as HC that in those identified as MC and LC (Figure 3B and Table S4). Considering that HC-lncRNAs were the most reliably predicted and the majoritarian class in all the species analyzed, further analyses were restricted to this lncRNA category. A more exhaustive analysis of HC-lncRNA by considering individually lincRNAs, NAT-lncRNAs, int-lncRNAs and SOT-lncRNAs reveals that regarding the GC content, the four types of lncRNAs showed values significantly inferior to those observed in PC genes (median of 43.22%), but superior to Intergenic regions (median of 34.80%) (Figure 3C). Among the four types, generally, NAT-lncRNAs (median of 40.84%) and SOT-lncRNAs (median of 39.73%) have higher GC contents than lincRNAs (median of 37.61%) and int-lncRNAs (median of 37.12%). Similarly, the exon number of the four lncRNAs types was significantly lower than that of PC genes (median of 4) (Figure 3C and Table S4). SOT-lncRNAs always showed more than 1 exon in all their transcripts in contrast to the other classes of lncRNAs, being the only one with an average of more than 2 exons (mean of 2.41).

It is worthy to note that in general, the average length of SOT-lncRNAs (median of 722 bp) and NAT-lncRNAs (median of 623 bp) was higher than the others types of lncRNAs (median of 355 and 352 bp, for lincRNAs and int-lncRNAs, respectively), although not as much as PC genes (median of 1275 bp). Consistently with other plant species, the expression level of the four types of cucurbits lncRNAs (median of 0.21, 0.43, 0.64 and 1.21 TPM, for lincRNAs, NAT-lncRNAs, int-lncRNAs and SOT-lncRNAs, respectively) was significantly lower than the observed for PC genes (median of 8.21 TPM) (Figure 3C and Table S4). When the repeat content was analyzed we observed that lincRNAs and int-lncRNAs have higher contents than PC genes surpassing, in some cases, the calculated value for random intergenic regions. These differences regarding to PC genes of GC content, exon number, length, expression and repeat content were consistently observed for all analyzed cucurbits (Figure S4).

### Evolutionary conservation of lncRNAs

We used sequence similarity and synteny as parameters to analyze the potential relatedness and to explore the evolutionary aspects of the HC-lncRNAs predicted in the nine cucurbit species. To infer sequence conservation, a BLAST pairwise alignment between species was performed with all the predicted HC-lncRNAs. Then, putative lncRNA orthologous families were identified by Markov Cluster (MCL) algorithm using the reciprocal best hits (RBH) from BLAST results. Additionally, a syntenic approach (36) based on 1:1 high-quality orthologous PC genes was employed to classify all the potential HC-lncRNAs into clusters referred hereafter as syntenic families. These families include lncRNAs from the different species that share the same genomic context, meaning that they are surrounded by orthologous PC genes. Considering as conserved the lncRNAs identified in at least two species, we observed that the positional conservation or synteny (mean of 50.97%) was significantly higher than the sequence conservation (mean of 35.41%) (Figure 4A and Table S5). In addition, as expected, the percentage of lncRNAs exhibiting primary sequence conservation was much lower than in PC genes (Figure S5A).

**Figure 4.**
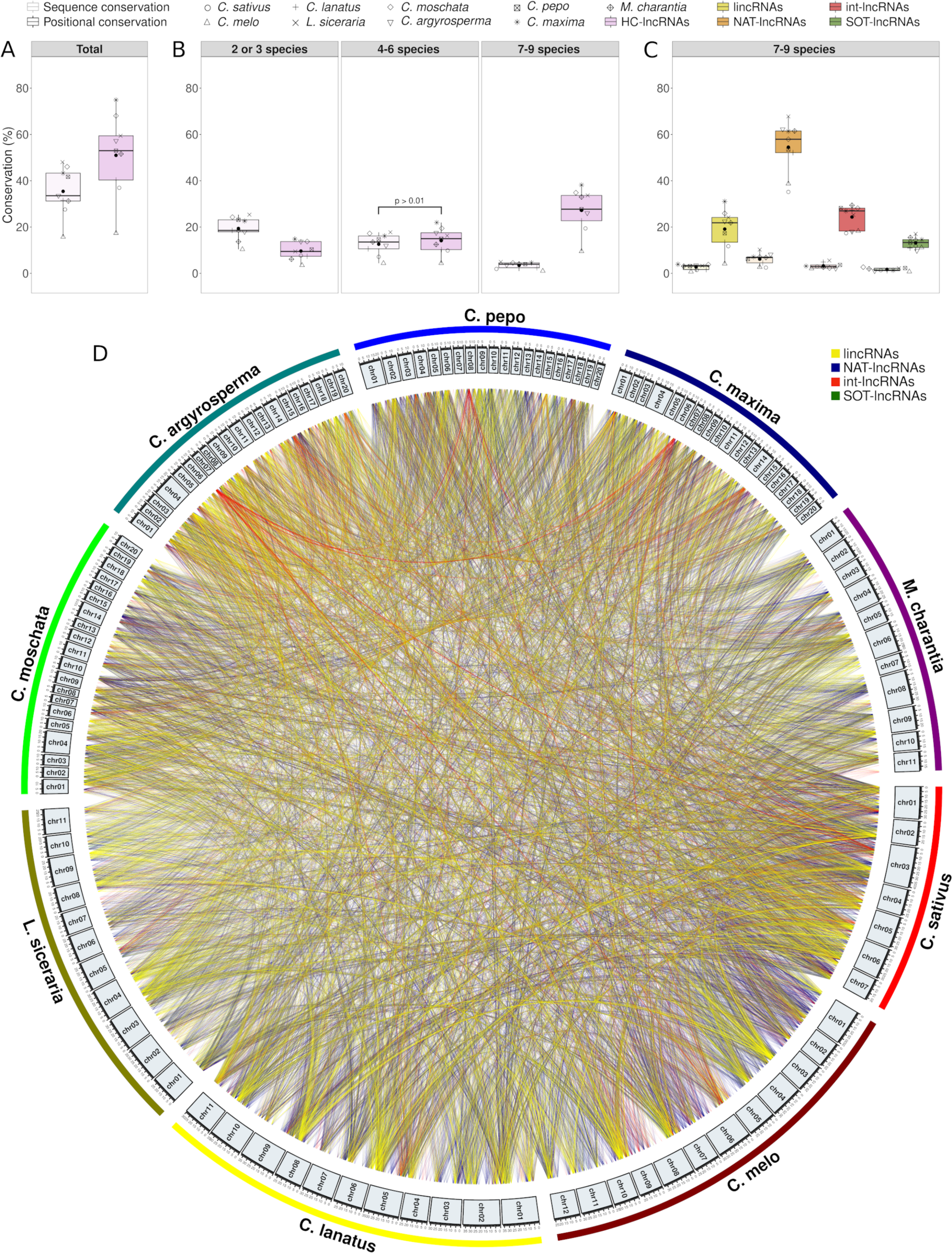
Conservation of the lncRNAs in the family *Cucurbitaceae*. (A) Box plots showing the percentage of lncRNAs identified as conserved at sequence (light color) and positional (dark color) levels. (B) Detailed information considering if lncRNAs are shared in two or three (left), four to six (center) and seven to nine species (right). (C) Values obtained for highly conserved lncRNAs (shared by 7 to 9 species) according to lncRNA classes (lincRNAs, NAT-lncRNAs, int-lncRNAs and SOT-lncRNAs). The nine species and the four-lncRNA classes are represented by different shape and colors, respectively (detailed on the top). Differences between pairs of box plots, sequence vs positional conservation, are statistically significant (Wilcoxon signed rank test p-value < 0.01), unless indicated otherwise. A black point represents the mean value. (D) Circos plot of the relationships of the conserved lncRNAs at the positional level (syntenic relationships). Each lncRNA class (lincRNAs, NAT-lncRNAs, int-lncRNAs and SOT-lncRNAs) is represented by a different color (detailed at the top right). In both sections, only lncRNAs identified as High-Confidence (HC-lncRNAs) were considered. Intergenic (lincRNAs), natural antisense (NAT-lncRNAs), intronic (int-lncRNAs) and sense overlapping (SOT-lncRNAs) lncRNAs.

Next, we compared the proportion of lncRNAs included in 3 categories of conservation: low (conserved in 2-3 species), medium (in 4-6 species) and high (in 7-9 species). The obtained results revealed that approximately 27.23% of the lncRNAs predicted in cucurbits were identified as highly conserved at the positional level, while sequence conservation in this category was lower than 5.0% (Figure 4B and Table S5). Further analysis showed that NAT-lncRNAs were the predominant class (mean of 54.48%) of highly positionally conserved lncRNAs and that conservation values obtained for lincRNAs, int-lncRNAs and SOT-lncRNAs were each less than 25% (Figure 4C and Table S5). Moreover, as in the comparison between sequence and positional conservation, the differences in sequence conservation between PC genes and lncRNAs identified as highly conserved were much greater (Figure S5B). Regarding the relation between the different cucurbit species, we observed that the syntenic relationships were comparable among all the species analyzed, except for *M. charantia* that showed significantly lower synteny levels. Positionally conserved lncRNAs were, in general, regularly distributed throughout the genome of the analyzed cucurbits (Figure 4D). Moreover, as expected from their closer evolutionary relationships, the majority of the syntenic relationships in lncRNAs with low and medium conservation were detected between species of the same genus (Figure S6 and S7, respectively). Finally, we analyzed transcripts for the presence of conserved sequence motifs shared by lncRNAs within a syntenic families and their statistical significance was calculated by generating datasets of randomized syntenic families (see Material and Methods for more details). We identify 982 syntenic families (30.13%) with at least one shared conserved motif. This proportion of syntenic families was significantly higher than what could be expected by chance as assessed in the simulated data (mean of 6.93%) (Figure S8A upper panel). Although comparable results were observed when each class of lncRNAs was considered individually, the higher differences in shared conserved motifs were detected in SOT-lncRNAs and int-lncRNAs (Figure S8A lower panel). We also compared the length (in nucleotides) and the statistical robustness (estimated by considering e-values) of the predicted motifs. The results showed that motifs identified in syntenic families were significantly longer and more reliable than those predicted in the simulated data (upper panels in Figure S8B and S8C, respectively). These significant differences were also observed when the different classes of lncRNAs were considered individually (Figure S8B and S8C lower panels).

### Expression of cucurbit lncRNAs is tissue-specific, development-dependent and responsive to environmental changes

To explore the potential biological properties of the predicted lncRNAs, we specifically analyzed projects included in our dataset in which different plant-tissues, developmental phases and/or changing environmental conditions have been considered. According to the criteria detailed in Material and Methods, 13 projects comprising eight cucurbit species were selected to analyze the tissue-specific expression of HC-lncRNAs and PC genes (see Table S6 and S7 for detailed information). Tissue-specificity was estimated using the TAU value which varies from 0 to 1, where 0 means broadly expressed, and 1 is specific (37). Our results showed that the tissue-specific expression ratio (TAU values) observed for the totality of the cucurbits lncRNAs (mean TAU 0.60), was significantly higher that the obtained for protein coding genes (mean TAU 0.37) (Figure 5A, left panel). These significant differences in the mean TAU values were also observed when each lncRNA class was considered individually. LincRNAs (mean of TAU 0.63) and NAT-lncRNAs (mean of TAU 0.60) were in general more specific in their expression in comparison to int-lncRNAs and SOT-lncRNAs, with TAU ratios of 0.56 and 0.49, respectively (Figure 5A, right panel).

**Figure 5.**
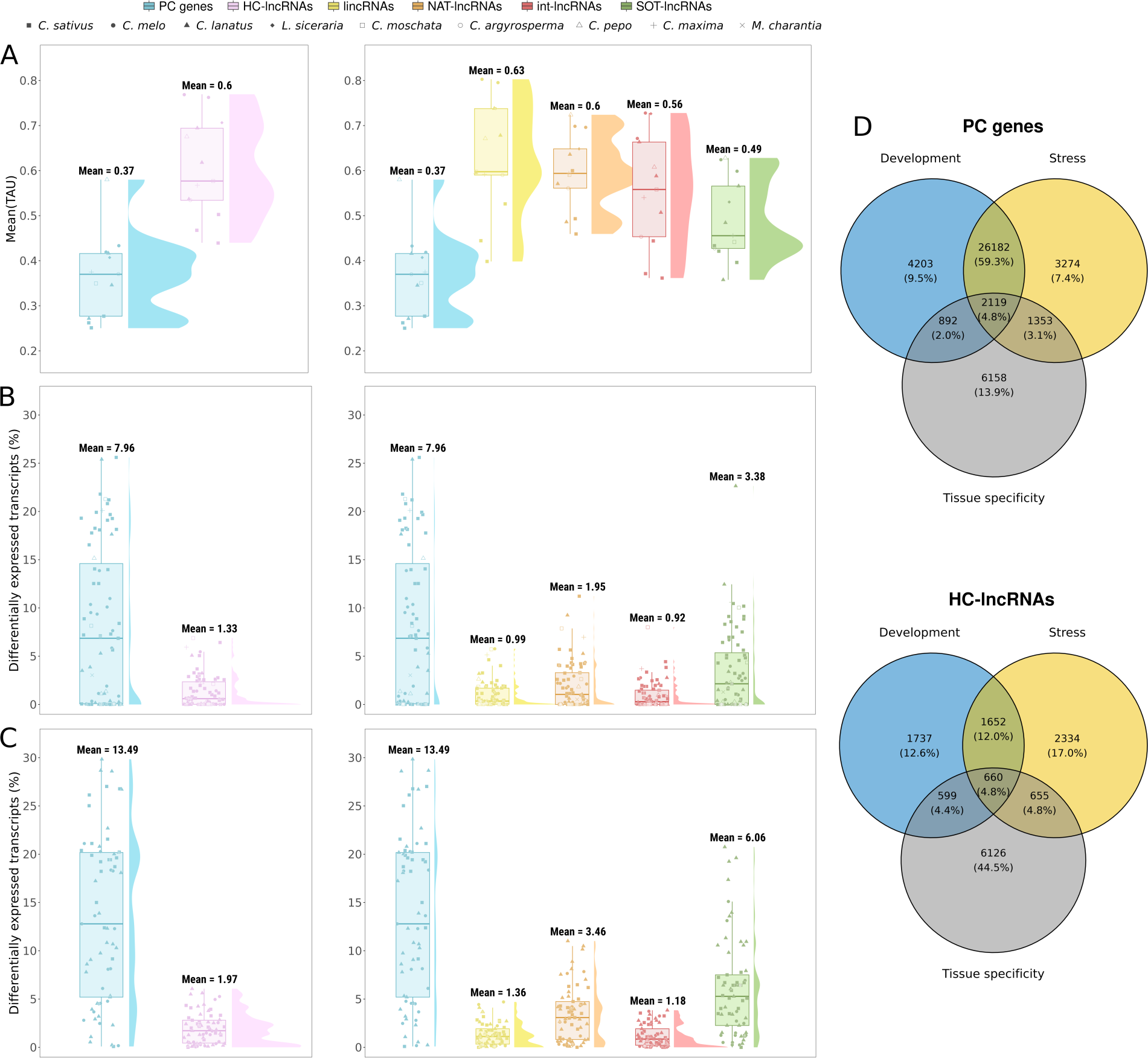
Expression of the lncRNAs considering different tissues, developmental stages and stress conditions. The boxplots in the left part represent the comparison between protein-coding genes (PC genes) and High-Confidence (HC-lncRNAs), while those in the right the comparison between PC genes and the four classes of identified lncRNAs: intergenic (lincRNAs), natural antisense (NAT-lncRNAs), intronic (int-lncRNAs) and sense overlapping (SOT-lncRNAs). (A) Comparison of the tissue-specificity calculated as the mean TAU value (0 means broadly expressed, and 1 is the most specific) (B) Comparison of the number of differentially expressed (DE) transcripts in response to stress conditions. (C) Comparison of the number of differentially expressed (DE) transcripts in different developmental stages. The nine species and the four-lncRNA classes are represented by different shape and colors, respectively (detailed on the top). (D) Venn diagrams showing the overlap between tissue-specific transcripts (TAU above 0.8) and those differentially expressed in specific developmental stages or in response to stress conditions. The upper diagram compares protein coding (PC) transcripts while the lower one lncRNAs. Differences between pairs of box plots, lncRNAs vs PC genes, are statistically significant (Wilcoxon signed rank test p-value < 0.01), unless indicated otherwise.

The correlation between different developmental stages and the expression of the predicted lncRNAs was analyzed in three cucurbit species comprising a total of 65 developmental events in 13 projects (see Table S6 and S7 for detailed information). The obtained results showed that the proportion (estimated as the mean value for the total of the studies) of lncRNAs with expression associated with development (1.97%) was significantly lower than the obtained (13.49%) when protein-coding genes were considered (Figure 5B, left panel). A more detailed analysis showed that SOT-lncRNA was the class with higher development-associated expression (Figure 5B, right panel).

Regarding environment-dependent expression we analyzed 75 stress events (with both biotic and abiotic origin) performed in seven cucurbit species (see Table S6 and S7 for detailed information). Our analysis demonstrated that the mean of the relative ratio of stress-responsive lncRNAs in cucurbits (1.33%), was also significantly lower that the observed for PC genes (7.96%) in the same studies (Figure 5C, left panel). In coincidence with the observed in development, transcripts classified as SOT-lncRNA were the predominant type of stress-responsive lncRNA (Figure 5C, right panel)

Finally, we selected the cucurbit species (*C. lanatus, C. sativus and C. melo*) with available data in the three selected conditions, to compare the specificity in the expression. Results showed in the Figure 5D demonstrate that predicted lncRNAs show (in comparison to PC genes) higher specific expression rate (defined as the percentage of specific transcripts) in the three analyzed conditions: tissue (44.5% vs 13.9%), stress (17.0% vs 7.4%) and development (12.6% vs 9.5%) (Figure 5D). These differences in specificity were also observed when each one of the three species was analyzed individually (Figure S9).

## Discussion

The tremendous achievement made over the last years in the computational prediction and functional biology of plant lncRNA has prompted the creation of detailed datasets containing description and information about this emerging type of regulators in diverse model and non-model species (8). However, the lack of systematic analyses performed from a more global point of view has meant that a significant number of economically relevant crops and closely related species can be poorly represented in these repositories. In this sense, members included in the *Cucurbitaceae* family, constitute a paradigmatic case of worldwide relevant crops, in which global information about lncRNAs is limited. Here, we have analyzed a large amount of available transcriptomic studies performed in nine representative species, to generate a comprehensive inventory of lncRNAs in cucurbits.

Importantly, our analysis is not restricted to lincRNAs as it happens in most studies of different plant species (8, 38). The use of directional RNA-seq data enabled the prediction of the other lncRNA categories: NAT-lncRNAs, int-lncRNAs and SOT-lncRNAs. This is especially relevant since the two latter categories are considerably understudied in plants, while NAT-lncRNAs have been associated with important physiological roles by regulating the expression of their cognate sense genes by different mechanisms (39–41), but their annotation in cucurbits was lacking.

To provide robustness to our inventory we established three confidence levels for the prediction: High (HC), Medium (MC) and Low (LC). Results obtained during our study revealed that HC-lncRNAs (beside to be the predominant) showed the highest structural and/or potential biological differences with PC genes. Consequently the majority (except when the contrary is specified) of the analyses showed in this work were performed with this lncRNA type. In coincidence with the observed in other plants, most of the lncRNAs predicted in the nine species, are derived from intergenic regions. Considering the amount of lncRNAs identified in each species, *C. melo*, *C. sativus* and *C. lanatus* have the largest number of predicted lncRNAs while *C. maxima*, *M. charantia* and *L. siceraria* the smallest. Although diverse reasons could be associated whit this difference, it is important to note that both groups are constituted by the species with the highest and lowest numbers of analyzed samples, respectively. In this sense, it has been proposed that the achievement of multiple transcriptome analysis under diverse environmental and physiological conditions in specific cell types can contribute to the discovery of more functional lncRNAs (3, 42).

The analysis of the main molecular features of the predicted lncRNAs revealed significant differences between these regulatory transcripts and PC genes and/or a set of randomly generated intergenic regions. Consistent with observations in other plants (7, 10, 43, 44), lncRNAs were shorter and have lower GC content and exon number than PC genes. In addition, and as expected, cucurbit lncRNAs also showed low expression levels. In this sense, a recent study support that, in mice, lower expression can be essential for the functional role of lncRNAs by ensuring specific recognition of its regulated targets, suggesting that low accumulation may be an essential feature of how lncRNAs work (45).

Another distinctive characteristic of the predicted lncRNAs was their high content of repetitive regions in comparison to PC genes, but significant smaller that in random intergenic regions. Interestingly, increased level of repetitive regions respect to PC genes has been described as a common characteristic for lncRNAs in several organisms (3). In general, these significant differences with PC genes tend to be more evident in lincRNAs, suggesting the existence of certain differential characteristics for this type of lncRNAs, may be related to their relevant regulatory role (42, 46, 47).

Considering the evolutionary relationships of the predicted lncRNAs, lower sequence conservation was observed in comparison with the positional conservation of the syntenic families. This divergence in conservation (higher at positional than sequence level) was increased when lncRNAs with high conservation rate (in almost 7 species). This evidences that during the evolution of cucurbits lncRNAs have mostly diverged in their sequences, which could be related to an increased phenotypic plasticity. Interestingly, syntenic lncRNAs also showed a significant tendency to contain more conserved and longer sequence-motifs than randomly generated transcripts. Overall, these results suggest that despite large evolutionary distances (evidenced by low sequence conservation), the analyzed cucurbits possess syntenic lncRNAs that share relatively conserved sequence-motifs that can be assumed like modules. This observation, is in consonance with previous studies (36, 48, 49) supporting that in general, lncRNAs tend to acquire a modular structure and are rich in repeats, implying that small sequence elements can be also key determinants of lncRNA function (50).

It is widely accepted that lncRNAs exhibit distinctive biological features such as tissue-specific expression, differential temporal regulation and expression related to environmental changes (3, 10, 43, 51, 52). The detailed analysis of our dataset demonstrated that lncRNAs predicted in cucurbits fulfilled these functional conditions. The obtained results revealed that lncRNAs identified in the eight analyzed species (having studies with multiple tissues) showed significantly higher TAU values those PC genes, supporting that their expression is mostly tissue dependent, in accordance with the observed for other plant lncRNAs (43, 44). The proportion of lncRNAs with differential expression was lower that obtained for PC genes. However, these differential lncRNAs showed (when compared with PC genes) a specific response pattern highly dependent of environment and development (see figure 5D). Interestingly, this high specificity in the expression pattern was also observed when tissue-dependent lncRNAs were analyzed. However, it cannot be excluded that, at least partly, this apparent specificity in the expression of lncRNAs could also be attributed to the generally low expression level of lncRNAs (43).

Both the differential structural and functional features support that the lncRNAs inferred here constitute a subset of RNAs significantly distinctive of the protein coding and/or randomly generated transcriptome. Therefore, in this study we provide a comprehensive catalogue of the different types of lncRNAs in the family *Cucurbitaceae* describing the confidence of their prediction, their molecular features and conservation (Table S8). Although we recognize that this is only a initial step and that additional experimental approaches and cucurbits species should be considered in the future, it is expected that this extensive inventory will constitute a valuable resource for further research lines focused on elucidate the basis of the regulation mediated by lncRNAs in cucurbits.

## MATERIAL AND METHODS

### Data preprocessing and assembly

With the aim to perform a comprehensive study, we selected all cucurbit species with fully sequenced genome and annotation file available (at 24th of January 2022) in the Cucurbit Genomics Database (CuGenDB) (32). Then, all RNA-Seq data publicly available (at 24th of January 2022) at the Sequence Read Archive database (SRA) (53), were retrieved using the commands prefetch and fasterq-dump provided by SRA Toolkit v.2.11.2 (https://github.com/ncbi/sra-tools). Only species comprising initially more than five RNA-seq samples were retained for quality control filtering and strandedness identification: *C. sativus, C. melo, C. lanatus, L. siceraria, B. hispida, C. moschata, C. argyrosperma, C. pepo, C. maxima, L. aegyptiaca, M. charantia* and *S. grosvenorii*. First, we removed adapters and filtered reads by quality (average quality > 20, window size = 4 bp) and length (length > 49 bp) using fastp v.0.23.2 (54) to provide clean data for downstream analysis. FastQC v.0.11.9 (https://www.bioinformatics.babraham.ac.uk/projects/fastqc/) and Multiqc v.1.11 (55) were used to perform quality control of clean data. Second, we identified the strandedness of the data and discarded non-strand-specific or non-stranded RNA-seq samples. Strand-specific or stranded RNA-seq samples provide a more accurate estimate of transcript expression and allow the identification of more types of lncRNAs such as natural antisense lncRNAs. To perform this step, we ran rsem-prepare-reference from the software package RSEM v.1.3.3 (56) to extract the transcriptomes of the analyzed species, and subsequently executed the pseudoaligner salmon v.1.6.0 (-l A) (57) in mapping-based mode, which identifies the strandedness of the data. Then, only species comprising more than five cleaned and strand-specific RNA-seq samples were included in the subsequent analysis.

Once the species and their samples were preprocessed and selected, we used the splice-aware sequence alignment program HISAT2 v2.2.1 (58) for mapping the clean data to the corresponding reference genomes. In this step, strandedness information was considered (--rna-strandness <STRANDEDNESS salmon code>) and maximum intron length was set to 10000 bp (--max-intronlen 10000), which corresponds to approximate maximum intron length in plant genomes. Moreover, --dta parameter was used to help transcript assemblers significantly improve computation and memory usage. Next, we reconstructed the transcriptome of each sample individually using the StringTie2 v2.2.0 assembler (59). In particular, we performed a genome-guided assembly approach taking into account the strandedness information (--rf/-- fr). All individual assembly gtf files produced by StringTie2 in the previous step were merged into a single and unified transcriptome for each species using the merge option of StringTie2 (-g 50 -F 0.3 -T 0). Finally, we compared the transcriptomes to the reference annotation files of each species to identify novel transcripts using the software GffCompare v.0.12.6 (60). According to their genomic location and referring to the neighboring protein-coding (PC) genes, this software classified the assembled transcripts and assigned them a class code.

### Computational prediction of lncRNAs

Based on the class code annotation, we selected transcripts annotated as “u” (intergenic), “x” (antisense), “i” (intronic) and, “o” or “e” (sense). Specifically, class code “u” refers to transcripts that come from intergenic regions of both genomic strands, class code “x” refers to transcripts that overlap with the exons of a PC gene on the opposite genomic strand, class code “i” refers to transcripts that are completely contained within the intron of a PC gene on the same genomic strand and class codes “o” or “e” refer to transcripts that overlap with the exons of a PC gene on the same genomic strand. Next, transcripts were filtered by length (length > 200 bp), exon number (exon number >= 1) and expression level (Fragments Per Kilobase Million > 0.3).

Further, we assessed the coding potential of the predicted transcripts using three alignment-free computational tools CPC2 v.1.0.1 (33), FEELnc v.0.2 (34) and CPAT v.3.0.2 (35) based on various intrinsic properties. For CPC2, transcripts assigned with the “noncoding” label were considered non-coding transcripts. For FEELnc, two training datasets, known PC genes and known lncRNAs, were required, and the shuffled mode was used to generate the later. The FEELnc cutoff to consider a transcript as coding or non-coding was defined using a tenfold cross-validation. For CPAT, we used the *Arabidopsis thaliana* logit model and hexamer frequency table obtained from the CREMA tool (61). The CPAT cutoff to consider a transcript as coding or non-coding was the default CREMA cutoff (0.5). To improve the robustness of the results, homologies were searched (on 18th of February 2022) with Swissprot (62) and Pfam-A databases (on 29th of May 2022) (63). On the one hand, we ran the blastx command (--strand plus --more-sensitive --top 5 --evalue 1e-5) from the program DIAMOND v.2.0.14 (64) to align the predicted transcripts to the protein sequences present in Swissprot. On the other hand, the TransDecoder.LongOrfs script (-m 20 -S) from the program transdecoder v.5.5.0 (https://github.com/TransDecoder/TransDecoder) was run to extract the longest Open Reading Frame (ORF) of each transcript and after that, the hmmsearch command (-E 1e-5 --domE 1e-5) from the software package HMMER v.2.0.14 (65) was used to align these longest ORFs to the protein domains present in Pfam-A database. Then, transcripts were classified into three confidence-levels according to the fulfillment of the following criteria: i) High (HC): predicted as lncRNA by the three software (CPC2, CPAT and FEELnc) and lacking homology with ORFs in the two protein databases (SwissProt and Pfam-A), ii) Medium (MC): predicted as lncRNA by the three software but have similarity to one protein database or lack similarity to the two protein databases but are only predicted by two software and iii) Low (LC): predicted as lncRNA by two software and have similarity to at least one protein database or lack similarity to the two protein databases but are only predicted by one software. Those transcripts that don’t meet any of the present scenarios were not classified.

The next step of the procedure was performed to filter out housekeeping non-coding RNA classes such as ribosomal RNA (rRNA), transport RNA (tRNA), small nuclear RNA (snRNA), small nucleolar RNA (snoRNA) and miRNA precursors (pre-miRNAs). To remove these housekeeping ncRNAs, we ran the blastn command (-strand plus -evalue 1e-5) from the NCBI-BLAST software v.2.13.0+ (66) to align the remaining transcripts from the confidence-level classification step to RNAcentral (on 15th of June 2022) (67). Only ncRNAs from the *Cucurbitaceae* family were considered in the alignment. To remove pre-miRNAs, we repeated the previous step against miRBAse (on 1st of June 2022) (68) and PmiREN (on 15th of June 2022) (69) using only pre-miRNAs from the analyzed species. Subsequently, significant hits were aligned to miRNA sequences from the analyzed species and from miRBase and PmiREN using blastn (-dust no -evalue 0.05 -word_size 7 -strand plus). Then, the program MIReNA v.2.0 (70) was used to validate potential pre-miRNAs with information on potential miRNAs. In order to provide additional information, we annotated filtered transcripts using potential known lncRNAs databases such as CANTATAdb v.2.0 (31), PLncDB v.2.0 (30) and GreeNC v.2.0 (29), last access on 15th of June 2022. Only lncRNAs from the analyzed species were downloaded and the program used to align the filtered transcripts to the different lncRNAs databases was blastn (-strand plus -evalue 1e-5). Only significant hits covering more than 50% of both aligned sequences were really considered hits. After that, when several isoforms were present, we kept only the longest one using the gtf2gtf command from the software CGAT v.1.0 (71). Finally, we created the database in table format as well as gtf annotation file format and the different classes of potential lncRNAs were renamed from intergenic, antisense, intronic and sense to lincRNA, NAT-lncRNA, int-lncRNA and SOT-lncRNA, respectively.

### Genomic distribution

Genome-wide distribution of potential lncRNAs and annotated PC genes was analyzed. Considering genomic density as the number of transcripts, we calculated the density per genomic window (size of 250 Kb) for the four classes of lncRNAs and PC genes using the R package circlize v.0.4.15. This information was displayed on a circular density plot using the same R package. After that, the percentage of the genome covered by each category (lncRNAs and PC genes) was obtained using bedtools genomecov v.2.27.1 (72). In addition, high-density regions (HDR) of the four classes of lncRNAs and PC genes were identified considering a window size of 100 Kb. In each case the threshold was defined as (i) the mean genomic density across all windows, (ii) multiplied by 1.5, and (iii) rounded up. All windows above their corresponding threshold were considered as HDRs. Finally, we calculated the percentage of HDRs of PC genes that overlaps with HDRs of each class of lncRNA.

### Molecular properties comparison

Potential lncRNAs and PC genes were compared using five features: GC content, exon number, length, expression level and repeat content. The first three features were obtained previously during the creation of the database. Expression level was calculated using the pseudoaligner salmon in mapping-based mode to quantify both transcript categories considering strandedness information (-l <STRANDEDNESS salmon code>). The relative transcript abundance was estimated in units of Transcripts Per Million (TPM), normalized by transcript length and library size. To analyze, the repeat content repeat calling was performed using RepeatModeler v.2.0.3 (73) and RepeatMasker v.4.1.3-p1 (73) on the cucurbit genomes. Then, bedtools intersect was used to calculate the proportion of each transcript in both categories covered by the repeat regions found. In addition, a third category of sequences corresponding to random intergenic genome regions (500 bp in length) was included to assess the GC content and repeat content. This new category was obtained using bedtools random (-l 500 -n 25000) on the cucurbit genomes and bedtools intersect to select those random sequences that don’t match any of the other two categories.

### Evolutionary conservation analysis

We assessed evolutionary relationships between predicted lncRNAs across species at sequence and positional level. To analyze the conservation at sequence level, pairwise alignments between species were performed using BLASTn. To do this, we built a custom BLAST database for the set of lncRNAs of each species. Then, each set of lncRNAs was aligned against each database with BLASTn (-strand plus) using a relatively non-stringent e-value threshold of 1e-5, and the best matched query-target pair was selected. After that, orthologous gene-pairs were identified based on Reciprocal Best Hits (RBH), and subsequently we ran OrthoFinder v.2.5.4 (74) based on the Markov Cluster (MCL) algorithm to infer the putative lncRNA orthologous families across species using RBHs information. To analyze the conservation at positional level, a syntenic approach developed and validated in (75) was implemented. This computational strategy identifies lncRNAs from different species that share the same genomic context, i.e. those surrounded by 1:1 orthologous PC genes. Therefore, we first ran OrthoFinder to infer high quality 1:1 orthologous PC genes in the analyzed species using the peptide sequences downloaded from CuGenDB. Next, pairwise comparisons between species were performed to search for syntenic relationships between lncRNAs. To this end, the code from (75) was automated so that we could compare as many species as we wanted to, and avoid any problem related to species names. Regarding the parameters, we kept the same as previously to compare the genomic context of lncRNAs between two different species: (i) Consider three transcripts on each side of a given lncRNA, (ii) a minimum of overall three shared PC genes (orthologs) and, (iii) a minimum of one shared PC gene on each side of a given lncRNA. Finally, pairwise syntenic lncRNAs were classified into clusters across species called syntenic families.

In addition, MEME v.5.5.1 from the MEME Suite (76) was used to identify conserved sequence motifs within the previously identified syntenic families. As parameters, we selected classic objective function, oops distribution, and a motif width between 6 and 50 nucleotides. Only motifs with e-value < 0.05 were considered as significant. As previously (36), we assessed the differences between the identified number of syntenic families with shared motifs and the random expectation. To do this, we randomly chose lncRNAs to generate 50 simulated datasets with the same number and size of lncRNA families as the real dataset and, preserving the number of lncRNAs per species observed in the real syntenic families. Then, we scanned the sequences of lncRNAs of these simulated families for motifs as described above.

### Tissue-specificity analysis

The tissue-specificity analysis was performed individually per RNA-seq experiment. Therefore, we first selected those experiments that had at least three different tissues in order to increase the robustness of the analysis. Next, we generated a table corresponding to the mean expression level (in TPM) of each potential lncRNA and PC gene per tissue. To be stringent, those transcripts that didn’t have more than 1 TPM in any of the tissues were filtered out. Finally, we used the python API of the tissue-specificity calculator tool tspex (https://github.com/apcamargo/tspex) to obtain the tissue-specificity metric Tau (77), which describes in a single value how tissue-specific or ubiquitous is a gene across all tissues, ranging from 0 (broadly expressed genes) to 1 (highly tissue-specific genes).

### Differential expression analysis in stress response and development

Given the regulatory role of lncRNAs in various developmental processes and stress responses (biotic and abiotic), all the projects present in this study related to both cases were selected for differential expression analysis (DEA). We first prepared the metadata to be able to carry out all pairwise comparisons (stress vs. control and late development stage vs. early development stage). Comparisons in which a condition had less than 2 replicates were discarded. Then, DEA was conducted by the R package DEseq2 (78), testing PC genes and potential lncRNAs. All p-values were adjusted by false discovery rate (FDR) and only transcripts with adjusted p-value ≤ 0.05 were considered as differentially expressed.

## CODE AND DATA AVAILABILITY

All necessary components including scripts, software versions and additional files needed to replicate the study findings can be found on our GitHub page (https://github.com/ncRNA-lab/Cucurbit_lncRNAs_landscape). The gtf annotation files of the identified lncRNAs are also available at this link. Accessions to the analyzed RNA-seq datasets are detailed as supplementary material.

## ACKNOWLEDGMENTS

This work was supported by the Agencia Estatal de Investigacion (AEI) (co-supported by FEDER - EU) Grants PID2022-1393930B-I00. The funders had no role in the experiment design, data analysis, decision to publish, or preparation of the manuscript.

## AUTHOR CONTRIBUTIONS

P.V.B. Searched downloaded and curated the data. P.V.B. Designed/adapted the pipeline and performed the bioinformatic analysis. P.V.B., J.M.M. and G.G. Designed the strategy, analyzed/discussed the results and wrote the paper. G.G. Conceived the general idea and drafted the manuscript. All authors read, revise and approved the final manuscript.

## CONFLICT OF INTEREST

All the authors declare no conflict interests.

## SUPPLEMENTARY FIGURES

**Figure S1.** Covered genome by protein coding genes and the predicted lncRNAs.

**Figure S2.** Genome distribution of predicted lncRNAs.

**Figure S3.** Molecular properties of all lncRNAs identified in each species.

**Figure S4.** Molecular properties of the four-lncRNA classes identified in each species.

**Figure S5:** Predicted lncRNAs exhibit lower sequence conservation.

**Figure S6:** Detailed information about syntenic relationships of lncRNAs positionally conserved in two to three species.

**Figure S7:** Detailed information about syntenic relationships of lncRNAs positionally conserved in four to six species.

**Figure S8.** Syntenic HC-lncRNAs exhibit modular structure.

**Figure S9.** Expression of the identified lncRNAs per species considering different tissues, developmental stages and stress conditions

## SUPPLEMENTARY TABLES

**Table S1:** Detailed description of the downloaded cucurbits dataset.

**Table S2:** Detailed description and global landscape of the lncRNAs predicted in cucurbits.

**Table S3:** Detailed information about the genome coverage of predicted lncRNAs.

**Table S4:** Detailed information about the molecular properties of the lncRNAs identified in cucurbits.

**Table S5:** Detailed information about the sequence and positional conservation values of the lncRNAs.

**Table S6:** Details of the dataset used for expression analysis in different tissues, developmental stages and stress conditions.

**Table S7:** Detailed information about lncRNAs expression associated to developmental stages, plant-tissues and response to stress.

**Table S8:** Detailed information about all the characteristics of the lncRNAs predicted in the nine species of cucurbits.

